# *Stomoxys* flies (Diptera, Muscidae) are competent vectors of multiple livestock hemopathogens

**DOI:** 10.1101/2024.10.07.611962

**Authors:** Julia W. Muita, Joel L. Bargul, JohnMark O. Makwatta, Ernest M. Ngatia, Simon K. Tawich, Daniel K. Masiga, Merid N. Getahun

**Author notes:** Corresponding author: Merid N. Getahun.

## Abstract

*Stomoxys* flies are widely distributed and economically significant vectors of various livestock pathogens of veterinary importance. However, the role of *Stomoxys* spp. in pathogen transmission is poorly understood. Therefore, we studied the feeding patterns of these blood feeders collected from specific locations in Kenya, to identify various vertebrate hosts they fed on, and the livestock hemopathogens they carried, to elucidate their role in pathogens transmission. Our findings show that field-collected *Stomoxys* flies carried several pathogens including *Trypanosoma* spp., *Anaplasma* spp., and *Theileria* spp. that were also found in the blood of sampled livestock, namely camels and cattle. The findings on blood meal analysis show that *Stomoxys* flies fed on a variety of domestic and wild vertebrate hosts. We further determined whether *Stomoxys* spp. are vectors of hemopathogens they harbored by studying the vector competence of *S. calcitrans, S. niger niger,* and *S. boueti* species complex, through laboratory and natural experimental *in vivo* studies. We show that in the process of blood feeding *Stomoxys* spp. complexes can transmit *T. evansi* (8.3%) and *T. vivax* (30%) to Swiss white mice. In addition, field-collected *Stomoxy* spp. were exposed to healthy mice for blood meal acquisition, and in the process of feeding, they transmitted *Theileria mutans* and *Anaplasma* spp. to Swiss white mice (100% infection in the test mice group). All mice infected with both trypanosomes via stomoxys bite died while those infected with *Theileria* and *Anaplasma* species did not, demonstrating virulence difference between pathogens. The key finding of this study showing broad feeding host range, cosmopolitan, plethora of pathogens harboured, and efficient vector competence in spreading multiple pathogens suggests profound role of *Stomoxys* on pathogen transmission and infection prevalence in livestock.

**Author summary:** *Stomoxys* flies are highly adaptable to several ecological settings, including metropolitan areas. In contrast, tsetse flies (genus *Glossina*), the main biological vectors of African trypanosomes, have a limited distribution to parks and other conservation areas. *Stomoxys* flies could play a significant role in the spread of animal African trypanosomes, among other hemopathogens, particularly in areas with or without tsetse infestation. Although there have been speculations about the potential role of *Stomoxys* flies in the transmission of various pathogens, there is lack of data to link hemopathogens occurring in both bloodmeal hosts of *Stomoxys* and in the flies, and further *in vivo* experimental studies to confirm the vector competence of Stomoxyine flies. Here, we explored a host and pathogens network, and investigated species diversity at various ecologies, and demonstrated that *Stomoxys* flies feed on diverse vertebrate hosts and are infected with a plethora of pathogens. We also showed experimentally that they could transmit some of these hemopathogens to mice, for instance, *T. vivax, T. evansi, Theileria mutans,* and *Anaplasma* spp. with varying infection success rates. *Stomoxys* flies could play a significant role in transmitting and spreading various hemopathogens of veterinary importance and possibly maintaining their circulation in livestock, which could explain the occurrence of animal African trypanosomes in the regions outside the tsetse belts.

## 1. Introduction

*Stomoxys* flies are widely distributed globally [1]. They feed on both blood [2], [3] and nectar [4]. These blood feeders pose a significant threat to livestock production worldwide, especially because of their occurrence in wider ecological zones [5]. The economic losses due to livestock infestation by *Stomoxys* flies are substantial, leading to a significant reduction in meat and milk production [6], due to pathogen infestation. These pathogens cause diseases including; animal trypanosomiasis, Rift Valley fever, African swine fever, lumpy skin disease, and anaplasmosis [3], [7–12]. The annual economic losses in the United States of America (USA) alone due to infestation by a single *Stomoxys* species, *Stomoxys calcitrans* – a cosmopolitan species commonly known as stable fly, have been estimated to be around US$ 2.2 billion [8]. The combined economic losses caused by *Stomoxys* flies infection, nuisance, and treatment costs outside the USA are not well documented at present.

*Stomoxys* feed on their vertebrate hosts’ blood once or twice a day [9]. Since the host animals respond to protect themselves against the painful blood-feeding fly bites, they (*Stomoxys*) typically do not complete blood-feeding on a single animal [5], [9]. In the event of an interrupted blood meal, the fly can restart feeding on another host by injecting infected saliva before feeding [5]. However, the pathogen transmission mechanism is not clear but could vary from biological, as is the case in the transmission of filariasis [10], [11], to the mechanical transmission of, for instance, animal trypanosomiasis [5], [12],[13].

Animal African trypanosomiasis caused by *T. vivax* and *T. evansi* is endemic in most countries in sub-Saharan Africa outside the tsetse belts [3], [12], [14], [15], and also in other regions outside Africa, including Asia [16], Latin America [17], and Europe [18]. *T. evansi* is the causative agent of ‘surra’, and *T. vivax*, that of nagana – an animal disease that is endemic in large swathes of Africa, Asia, and Latin America, and also present in the Canary Islands (Spain) [22] [23]. However, the establishment, widespread occurrence, maintenance, and circulation of *T. vivax* and *T. evansi* particularly in areas outside the tsetse belts is still unclear.

Despite the wide geographic distribution and cosmopolitan nature of *Stomoxys* flies, our knowledge about their blood meal hosts, and the hemopathogens they carry, which will provide insight into ‘vector-host-pathogen’ interactions, and disease transmission dynamics is limited and necessitates compressive research. Thus, in this study, we studied the diversity of *Stomoxys* spp. collected from various National Reserve to zero grazing ecologies (Fig 1). We conducted a study to assess the vector competence of *Stomoxys* spp., which refers to their host feeding dynamics, and capacity to become infected and transmit hemopathogens. We show that *Stomoxys* flies are competent vectors to multiple pathogens and may play a role in the transmission cycles of the pathogens they harbor.

**Figure 1.**
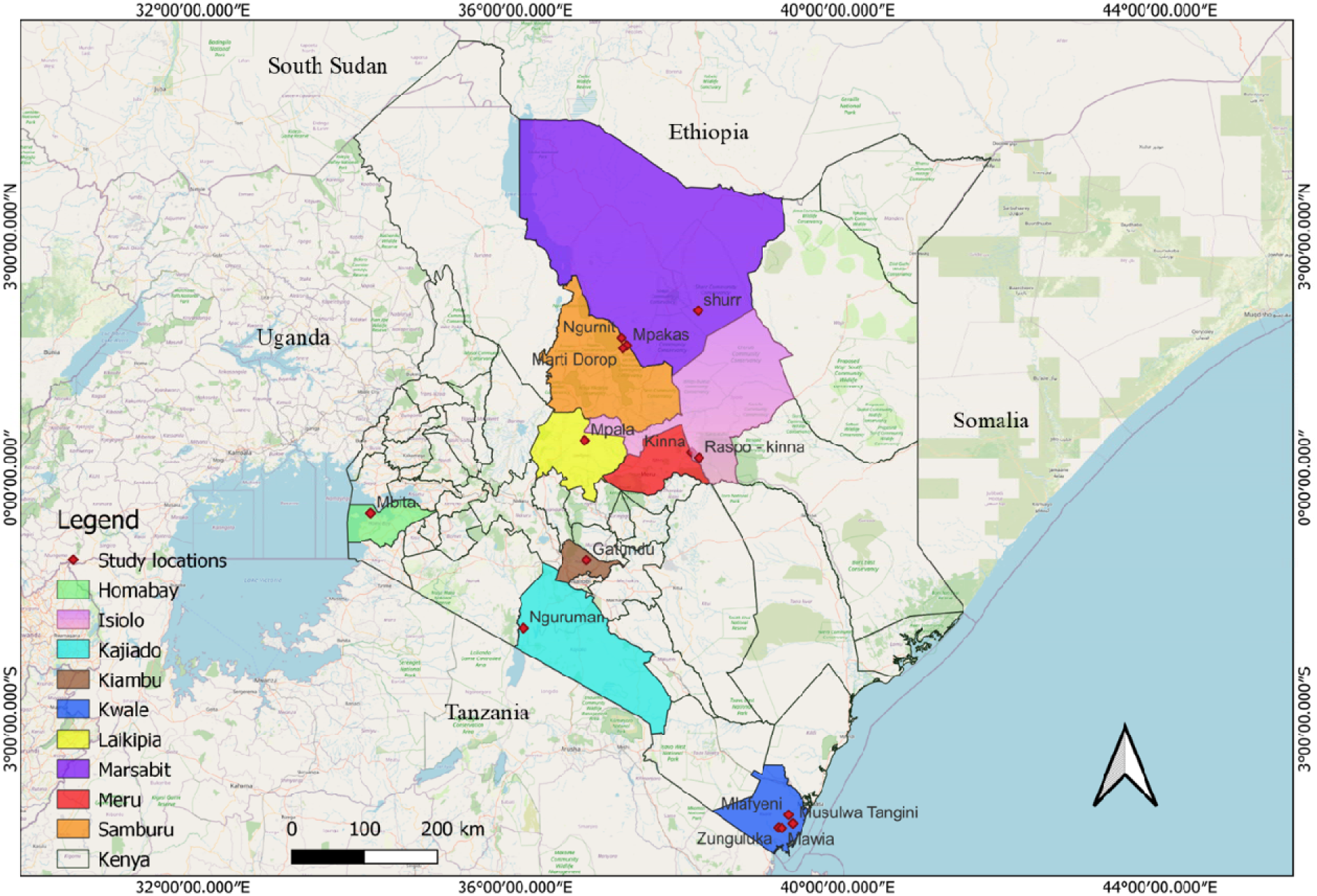
A map of Kenya showing the sampling sites across the nine selected counties. This map was created using the open-source software, QGIS v.3.

## 2. Materials and methods

### 2.1 Study sites

Field sampling took place in nine selected counties in Kenya at various times from October 2021 to November 2023. The sampled ecosystems in each County were inhabited by a variety of livestock species, wildlife hosts, and humans. These counties included: Kwale County (4.2572° S, 39.3856° E), Isiolo County (0.3257° N, 38.1961° E), Samburu County (1.7299° N, 37.3079° E), Kiambu County (1.0131° S, 36.9051° E), Kajiado County (1.7617° S, 36.0255° E), Marsabit County (2.4426° N, 37.9785° E) Homabay County (0.4368° S, 34.2060° E), Laikipia (0.2924° N, 36.8985° E), and Meru County (0.0515° N, 37.6456° E) (Fig 1).

### 2.2 Ethical approval and animal welfare

This study was conducted in strictly adherence to the approved experimental guidelines and procedures set forth by the Animal Care and Use Committee (IACUC) of the International Centre of Insect Physiology and Ecology, *icipe* (REF No.: icipeACUC2018-003-2023) and the Ethics Review Committee of Pwani University (REF No.: ERC/EXT/002/2020E). Farmers and pastoralists were sensitized about the research study and how the findings could benefit the farmers. Sample collection from livestock was done after obtaining verbal consent from farmers, as most of them (camel herders) were unable to read or write. Animal experiments complied with the approved guidelines and mice were handled carefully to ensure minimum distress.

### 2.3 Blood collection and fly trapping

#### 2.3.1 Blood collection

Blood collection from a total of 452 camels and 124 cattle was done from October 2021 to November 2023. The sample size was determined using our initial data, which indicated an infection rate of 4% in camels and 5.7% in cattle caused by *Rickettsiae* spp. the lowest prevalence among pathogens. We utilized this information as a basis for calculating the sample size, following the formula; 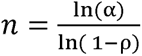 as outlined by [20]. At least 5 mL of blood was drawn from the jugular vein of each animal and collected in vacutainer tubes containing disodium salt of ethylene diamine tetra123 acetate (EDTA) (Plymouth PLG, UK). The blood samples were initially stored in vacutainers at 4°C until the collection was completed which were transferred to cryovials and stored in liquid nitrogen before being transported back to the Nairobi Duduville campus-*icipe* for molecular identification of pathogens.

#### 2.3.2 Trapping of flies

Flies were trapped using five red monoconical traps [21]. The traps were placed 150 meters apart and the flies were trapped from nine counties as indicated in (Fig 1). The traps were emptied twice per 24-hour period to avoid flies drying for clear identification and further intended analysis. The flies were later immobilized and preserved in absolute ethanol for further analysis and fresh flies were pinned for morphological identification. Trapping lasted for 5 days. Except for Kiambu County, all trapping sites were located in wide woodland savannah ecologies that were not in close proximity to villages.

### 2.4 Morphological identification of *Stomoxys* spp

Field-collected *Stomoxys* flies were taxonomically identified to the species level using established taxonomic keys as outlined by [22]. Briefly, the flies were staged under a dissecting Stemi 2000-C microscope (Zeiss, Oberkochen, Germany) to identify key morphological differences. The head frontal index and the distinctive dorso-abdominal patterns on the second and third segments were key distinguishing features in identifying fly sex and species, respectively. Images were captured using a digital microscope connected to an Axio-cam ERc 5s camera (Zeiss). The flies were grouped according to their species and preserved at −20°C for molecular analysis.

### 2.5 Molecular identification of *Stomoxys* spp. and livestock hemopathogens

#### 2.5.1 DNA extraction from blood samples and flies

In the initial stages of DNA pre-extraction, the flies were subjected to a 1% sodium hydroxide immersion for 1 minute to eliminate any exogenous material on their bodies. Subsequently, they underwent a 1-minute rinsing procedure with 1 × phosphate-buffered saline (pH = 7.4). Each fly was mechanically homogenized in sterilized 1.5-ml microfuge tubes containing 750 mg of 2.0-mm yttria-stabilised zirconium (YSZ) oxide beads (Glen Mills, Clifton, NJ, USA) and 80 µL of 1× PBS using a Mini-Beadbeater-16 (BioSpec, Bartlesville, OK, USA) for one minute. Genomic DNA extraction was done using DNeasy blood and tissue kit (Qiagen, Hilden, Germany) following the manufacturer’s protocol to obtain DNA from flies, cattle, and camel blood for pathogen screening. The obtained DNA was quantified using a Nanodrop spectrophotometer (Thermo Scientific, Wilmington, DE, United States) by comparing absorbance at 260 and 280 nm and later stored at −20°C for molecular work.

#### 2.5.2 Molecular identification of *Stomoxys* spp. and livestock hemopathogens

The extracted DNA was used in the molecular characterization of *Stomoxys* spp. as well as screening livestock hemopathogens. We conducted amplification using genus-specific primers as outlined in (Supplementary Table 1), using the conventional ProFlex PCR systems thermocycler (Applied Biosystems, Foster City, CA, USA). The PCRs were conducted in 10-μL reaction volumes, consisting of 5 µL nuclease-free water, 2 μL 5× HOT FIREPol® Blend Master Mix (Solis BioDyne, Tartu, Estonia), 0.5 μL of 10 µM forward and reverse primers (Supplementary Table 1), and of 2 μL DNA template. For negative controls, 2 μL nuclease-free water was used in place of the DNA template. The amplification conditions were set as described by [23]. PCR amplicons were resolved by electrophoresis under 2% ethidium-bromide-stained agarose gel (100 V for 1 hour), followed by DNA visualization by UV-transillumination (Kodak Gel Logic 200 Imaging System, CA, USA). PCR amplicons were purified using ExoSAP-IT (Affymetrix, Santa Clara, CA, USA) as per the manufacturer’s protocol, and outsourced for Sanger-sequencing at Macrogen Inc. (Amsterdam Netherlands).

#### 2.5.3 Phylogenetic analysis

To get a better understanding of how the various *Stomoxys* species and livestock hemopathogens are similar and different at the genomic level we performed a phylogenetic tree analysis. The nucleotide sequences acquired in this study were cross-referenced against the known sequences in the GenBank of NCBI nr database (https://www.ncbi.nlm.nih.gov/genbank/). BLAST was used to validate their identity and establish connections with existing deposited sequences [24]. To show the evolutionary relationships, maximum-likelihood phylogenetic trees were constructed using PhyML v. 3.0, employing automatic model selection based on the Akaike information criterion. The tree topologies were estimated through 1000 bootstrap replicates, incorporating nearest-neighbor interchange improvements [25]. The resulting phylogenetic trees were then visualized using FigTree v. 1.4.4[26].

#### 2.5.4 PCR-HRM vertebrate bloodmeal source identification in *Stomoxys* flies

The extracted DNA was used in the analysis of the blood meal source of the fed flies collected in the field. A 10-μL PCR reaction was carried out, consisting of 1 μL of DNA template, 6 μL of nuclease-free water, 2 μL of 5× HOT FIREpol EvaGreen HRM Mix from Solis BioDyne in Tartu, Estonia, and 0.5 μM of both forward and reverse primers (Supplementary Table 1). Positive control vertebrate host samples used as reference include; cow (*Bos taurus*), camel (*Camelus dromedarius*), donkey (*Equus asinus*), warthog (*Phacochoerus africanus*), African buffalo (*Syncerus caffer*), goat (*Capra aegagrus hircus)*, waterbuck (*Kobus ellipsiprymnus*) elephant (*Loxodonta africana*), sheep (*Ovis aries*), reticulated giraffe (*Giraffa reticulata*), lesser kudu (*Tragelaphus imberbis*), cheetah (*Acinonyx jubatus*), zebra (*Equus quagga*), baboon (*Papio*), gerenuk (*Litocranius walleri*), hartebeest (*Alcelaphus buselaphus*), reedbuck (*Redunca redunca*), hyena (*Crocuta crocuta*), gazelle (*Gazella gazella*), impala (*Aepyceros melampus*), lion (*Panthera leo*), bongo (*Tragelaphus eurycerus*) and human (*Homo sapiens*). The PCR thermal cycling conditions for primer were set as described by [3]. After PCR amplification, High-Resolution Melting (HRM) analysis was conducted within normalized temperature ranges, from 65°C to 78°C and 88°C to 95°C. The distinct melt curve profiles of the samples were compared against reference standards.

### 2.6 Experimental infection assays to determine the vector competence of *Stomoxys t*o transmit *T. evansi* and *T.vivax*

#### 2.6.1 Experimental animals

Swiss White Mice (*Mus musculus*) obtained from *icipe’s* Animal Rearing and Quarantine Unit (ARQU) were used for the infection experiment. Both male and female mice used for experiments were about 6 – 8 weeks old. Each mouse weighed about 24 – 29 g live body weight. The mice were housed under normal conditions in standard mouse cages and their diet primarily comprised of commercial pellets (Unga^®^ Kenya Ltd) and water, which was provided *ad libitum*. The mice were kept in a mice experimental room that was free from biting flies. The experimental mice used in pathogen transmission studies were not immune suppressed.

#### 2.6.2 Establishment of laboratory colonies of *Stomoxys* spp

*Stomoxys* spp. of both sexes were trapped from both Gatundu (1.0131° S, 36.9051° E) and around *icipe*-Duduville campus (1.2921° S, 36.8219° E) and taken to *icipe’s* insects rearing units. The mixed species of *Stomoxys* flies were maintained in 75 cm × 60 cm × 45 cm perspex cages (Astariglas^®^, Indonesia) and fed once daily between 9 am – 11 am warm defibrinated bovine blood obtained from a local slaughterhouse (Choice meats, Nairobi) and supplemented with 10% glucose and *Parthenium hysterophorus* flowers [27]. The temperature and humidity in the rearing room were kept at 25 ± 1 °C and RH 50 ± 5%, respectively with a 12:12 light/dark photoperiod. Sheep dung was used as an oviposition substrate [28] and developed pupae were picked and transferred to another cage for emergence. The newly emerged or teneral flies were used for experimental infection assays.

#### 2.6.3 Multiplication of *T. evansi* and *T. vivax* isolates in donor mice

The strain of *Trypanosoma* used in this study was *T. evansi,* which was isolated from a naturally infected camel (*Camelus dromedarius*) from Marsabit County, and *T. vivax* IL 2136 which was taken from *icipe’s* trypanosomes bio-bank. They were let to thaw after which parasitemia was checked using microscopy (Zeiss Primo Star Binocular Microscope, Zeiss, Oberkochen, Germany) with a ×40 magnification to ensure the viability of the stabilates before each inoculation. 200 µL of each stabilate was then inoculated to the mice through the intraperitoneal route

#### 2.6.4 Monitoring parasitemia levels in donor mice

The mice were monitored daily, three days after pathogen inoculation. Briefly, a drop of blood from the snipping of the mouse tail using a blood lancet was placed on a clean slide and covered using a coverslip as a wet blood smear and examined under a microscope (Seamer et al., 1993). The parasitemia score was estimated which correlated to a score sheet, as described by [29]. The period taken from the day post inoculation (dpi) to the first appearance of trypanosomes in blood was recorded for all mice. This was done until the required parasitemia was achieved (1 × 10^8^ trypanosome/mL blood).

#### 2.6.5 Determination of *T. evansi* and *T. vivax* survival rates in *Stomoxys* fly

Given that mechanical transmission of trypanosome parasitemia is known to be dose-dependent [30], our experimental design included the testing of two doses that are typically encountered in natural infection [3] approximately 1 × 10^8^ trypanosomes/ml blood and 5 × 10^8^ trypanosomes/mL blood. Following the successful induction of high parasitemia, the next step involved the extraction of whole blood from the donor mouse. This was done by sacrificing the mouse in accordance to the standard protocol defined by the Institutional Animal Care and Use Committee (IACUC). Fino-Ject disposable syringe 5 mL/cc with needle was used for blood collection by cardiac puncture [31] which resulted to 200 μL of blood. The parasitemia was checked microscopically, to ensure the blood still had enough parasite concentration. The infected blood was then carefully diluted in clean pre-warmed defibrinated bovine blood collected from the slaughterhouse (Choice meats, Nairobi) at a 1:1 ratio. The resulting blood mixture, approximately 400 µL, was applied onto clean cotton placed in a petri dish. A total of 60 teneral *Stomoxys* flies which were starved for 24 hours were placed in a clear acrylic plastic cage with dimensions of 10 × 10 × 15 cm, which was made of a 6-mm-thick perspex sheet from Astariglas® in Indonesia. The flies were fed on the blood-soaked cotton wool that was provided on a petri dish. Following a feeding period of five minutes, until all flies were fully engorged, the infection and spread of trypanosomes within the *Stomoxys* flies were monitored at various time points post-feeding, beginning one hour after the feeding event and continuing at each subsequent hour. To analyze the distribution and prevalence of the parasite within the bodies of the *Stomoxys* flies, we conducted dissections of various body parts, including the mouthparts, crop, and gut. At least five insects per exposure time were examined after immediate interrupted feeding.

#### 2.6.6. Experimental trials of in vivo transmission of *T. evansi* and *T. vivax*

Various protocols were tested in the *in vivo* transmission trials for optimization. At first, after the successful induction of high parasitemia within the donor mouse, the donor mouse and recipient mouse were restrained using a restrainer which was made of stainless-steel woven wire mesh with measurements of 0.9 mm per hole and a 400 µm wire diameter, and placed in a 10 ×10 ×15-cm cage made of 6-mm (thick) perspex clear acrylic plastic sheet (Astariglas®, Indonesia). The teneral flies n = 20 were introduced and the flies were disturbed by the observer to allow them to move from donor to recipient mice. Unfortunately, after more than 20 trials using this experimental method, we did not get any results. We optimized our experiment whereby, once a donor mouse with high parasitemia was achieved, the donor mouse and recipient mouse were restrained using a restrainer which was made of stainless-steel woven wire mesh with measurements of 0.9 mm per hole and a 400 µm wire diameter, and both were placed in separate 10 ×10 ×15-cm cage made of 6-mm (thick) perspex clear acrylic plastic sheet (Astariglas®, Indonesia). One-day-old teneral flies which were not fed on any blood meal where n=20 per experiment, were released in the cage of the donor mouse and allowed to feed for ≤1 minute. The timing was done once the proboscis had pierced the mouse’s skin to ensure feeding had started and the flies ingested blood from the infected mouse. This was followed by disrupted feeding where only the fed flies were individually picked using a respirator and transferred to the next cage containing restrained recipient healthy mice and flies were allowed to complete their blood meal until fully engorged (Fig 2). The flies were subsequently dissected to confirm the presence of parasites in their gut to assess the parasite-feeding success rate. The restrained recipient mouse was then released into their standard mouse cages and monitored daily after three days post-infection.

**Figure 2:**
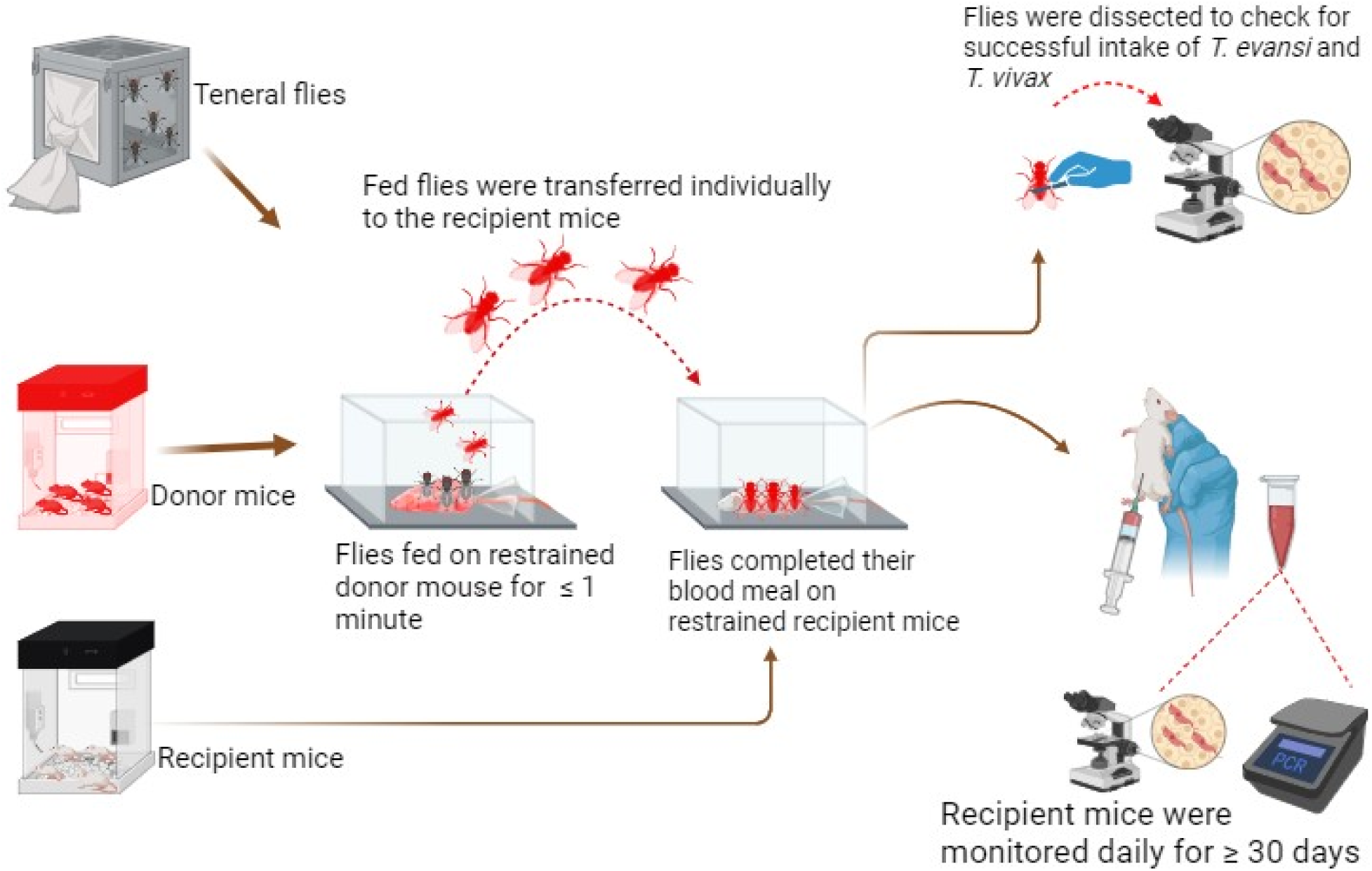
Laboratory *in vivo* transmission of *T. evansi* and *T. vivax* experimental design.

#### 2.6.7 Screening for *T. evansi* and *T. vivax* in the recipient mice by PCR

After three days post-infected fly bites a combination of microscopy and molecular methods were used to confirm the presence of parasites in the blood of infected animals for up to 30 days post-infection (dpi). For microscopy, it was done as described above, daily. Molecular screening was done by collecting blood samples from snipping the mice tails and collecting them in 1.5 ml eppendorf tubes which contained 80 μL 1× PBS buffer, pH = 7.4. Blood collection was done after every two days. This was followed by total DNA extraction using a DNeasy blood and tissue kit (Qiagen, Hilden, Germany) following the manufacturer’s protocol. PCR, gel electrophoresis, and gene sequencing were performed as described above

### 2.7 Determination of vector competence through field bioassay

Fly trapping was done as described above in various study sites. The traps were emptied after 6 hours and the flies were put in 10 × 10 × 15-cm cage made of 6-mm (thick) perspex clear acrylic plastic sheet (Astariglas^®^, Indonesia). The recipient mice were restrained using the restrainer that was made of stainless-steel woven wire mesh with measurements of 0.9 mm per hole and a 400 µm wire diameter and released into the cage. The flies were left to feed for 30 minutes before releasing the mice. This was followed by daily evaluation of pathogens in the recipient mice through microscopy and molecular screening as described above.

### 2.8 Data analysis

The Shannon diversity index (H) was utilized to define the diversity index of biting flies among study counties and was calculated using R statistical software (R version 4.4.1.). Estimated minimum infection rates (MIRs) of pathogens obtained for the flies were calculated as the number of positive per total number of flies tested ×100. Graphs were visualized using GraphPad software (GraphPad Software, Inc, USA). The *bipartite* R package’s interaction network [32] visualized the vectors blood-feeding behavior and pathogen interactions between hosts and vectors which was generated by R statistical software (R version 4.4.1.). An *Upset* plot displayed the number of flies feeding on specific animal species and those containing bloodmeals from one or more host species and was plotted using R statistical software (R version 4.4.1.). Transmission rates of the experimental infection assays were performed by calculating the number of infected mice per total number of transmission trials done ×100.

## 3. Results

### 3.1 *Stomoxys* species diversity relative abundance is ecology dependent

Diverse *Stomoxys* species were trapped throughout the year. A total of 11,323 adult *Stomoxys* flies were collected from various sites, from National Reserve including; Shimba Hills National Reserve and Nguruman Conservation Reserve, to zero grazing ecologies and identified morphologically using specific keys to the species level according to [22] as *S. calcitrans, S. sitiens, S. niger niger, S. niger bilineatus, S. boueti,* and *S. taeniatus* (Fig 3A). The sampling sites varied in species richness with some having only three species, while others had up to six. *S. calcitrans* was identified in all study sites while *S. taeniatus* was only found in Kajiado County. *S. calcitrans* exhibited a body appearance characterized by three dark spots on each of the second and third segments. *S. sitiens* abdominal segments resemble that of *S. calcitrans* but the dark spots are more transversely elongated. *S. niger niger* appears to have grey coloration with well-defined and dark stripes on abdominal segments. The dorsal view of *S. boueti* appears to have an indistinct dark abdomen and is much smaller in size. *S. niger bilineatus* has a brownish appearance with the abdominal segment having defined the dark stripes in a dorsal view. S*. taeniatus* has a brighter golden brown to almost yellowish color and it is larger than the other species (Fig 3A). Our molecular taxonomy using CO1 (Cytochrome Oxidase I gene) DNA sequence confirmed the morphological identification of those samples clustered distinctly and with previously documented DNA sequences demonstrating they are different species (Fig 3B). New CO1 sequences including *S. boueti,* (GenBank Accession number, PP587243) and S*. taeniatus* (GenBank Accession number PQ203543) were deposited that were not available in the NCBI.

**Figure 3:**
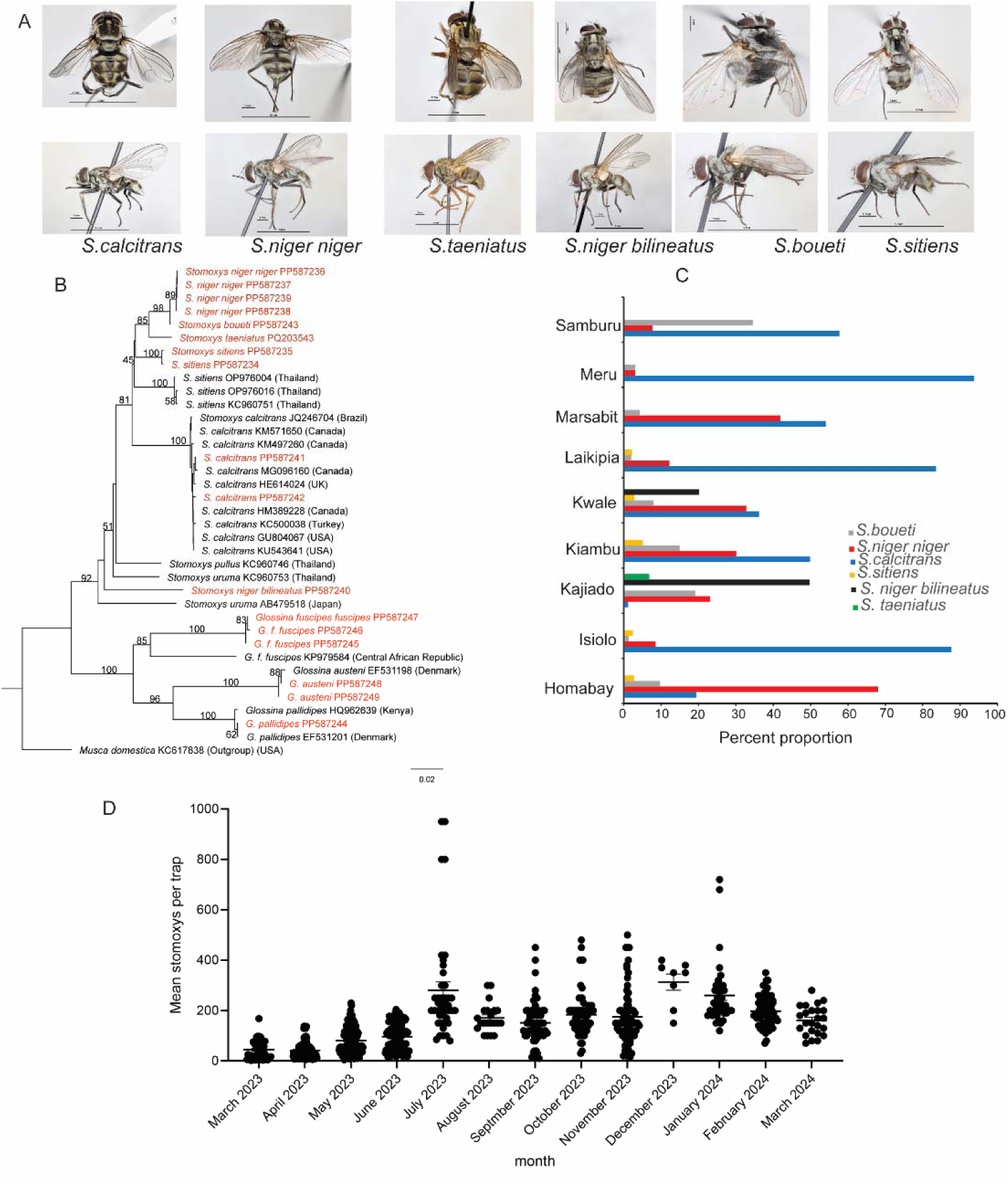
*Stomoxys* flies morphological identification, molecular characterization, species diversity, and seasonality (A) Image showing the dorsal and lateral view demonstrating the distinct morphological features of the six *Stomoxys* species encountered in various study sites (B) Neighbour-joining tree constructed based on aligned sequences of CO1 tree showing the relatedness of the various *Stomoxys* species. (C) The species diversity and their relative abundance in various ecologies. (D). Seasonality of *Stomoxys* at Gatundu site from Kiambu County.

Kajiado followed by Kwale County recorded the highest number of *Stomoxys* species diversity, six and five species, respectively (Fig 3C) The Shannon diversity index shows the varying levels of *Stomoxys* species diversity (Supplementary Table 2) across Kenyan counties with Kwale County having the highest Shannon diversity index of 1.36 and Meru County having the lowest Shannon diversity index of 0.29. Overall, the species distribution showed that *S. calcitrans* was the dominating species, except in Kajiado and Homabay counties, which was accounted for by (n= 5,547, 49%), followed by *S. niger niger,* (n= 2,938, 25.95%), *S. boueti* (n= 1,471, 12.99%), *S. niger bilineatus* (n= 778, 6.87%), *S. sitiens* (n= 495, 4.37%), and finally *S. taeniatus* (n=94, 0.83%). Using one of our sites (Kiambu County) we studied the seasonal dynamics of *Stomoxys* flies. *Stomoxys* flies were caught all year round with seasonal variation (Fig 3D). The abundance of *Stomoxys* increase with the rainfall data, there was an annual rainfall of 674 mm with a monthly average of 56 mm in the study County during the study period.

### 3.2 *Stomoxys* flies blood meal host network analysis demonstrates *Stomoxys* flies feed on a wide host range

*Stomoxys* flies feed on diverse wild and domestic animals. In total 225 fed *Stomoxys* flies were successfully identified to analyze the *Stomoxys*-host feeding network. Fifteen distinct vertebrate blood-meal hosts were identified, including cattle (*Bos taurus*), camel (*Camelus dromedarius*), warthog (*Phacochoerus africanus*), African buffalo (*Syncerus caffer*), goat (*Capra aegagrus hircus)*, waterbuck (*Kobus ellipsiprymnus*) elephant (*Loxodonta africana*), sheep (*Ovis aries*), reticulated giraffe (*Giraffa reticulata*), zebra (*Equus quagga*), baboon (*Papio*), reedbuck (*Redunca redunca*), gazelle (*Gazella gazella*), impala (*Aepyceros melampus*), and human (*Homo sapiens*) (Fig. 4). *S. calcitrans* had the most diverse blood meal hosts followed by *S. boueti* and lastly *S. niger niger* (Fig 4). We observed the diversity of blood meal sources is dependent. Wildlife conservation (Shimba Hills National Reserve) had the most variety of identified blood-meal hosts. Kiambu and Meru counties had the lowest host diversity due to zero grazing in regions where *Stomoxys* were trapped, resulting in a limited number of hosts mostly only cattle. In general, cattle were the most detected and most preferred host across all species *S. calcitrans* (n=65/265), *S. niger niger* (11/205), and *S. boueti* (n=13/205) (Fig.4). Multiple host feeding was also revealed in some flies where HRM melt curves revealed two peaks that matched the standard reference (Fig. 4). This was most commonly found in livestock, including cattle and goats, cattle and sheep, cattle and camels, and detected once in wildlife, including waterbuck and buffalo may be due to interrupted feeding before completion (Supplementary Table 3).

**Figure 4:**
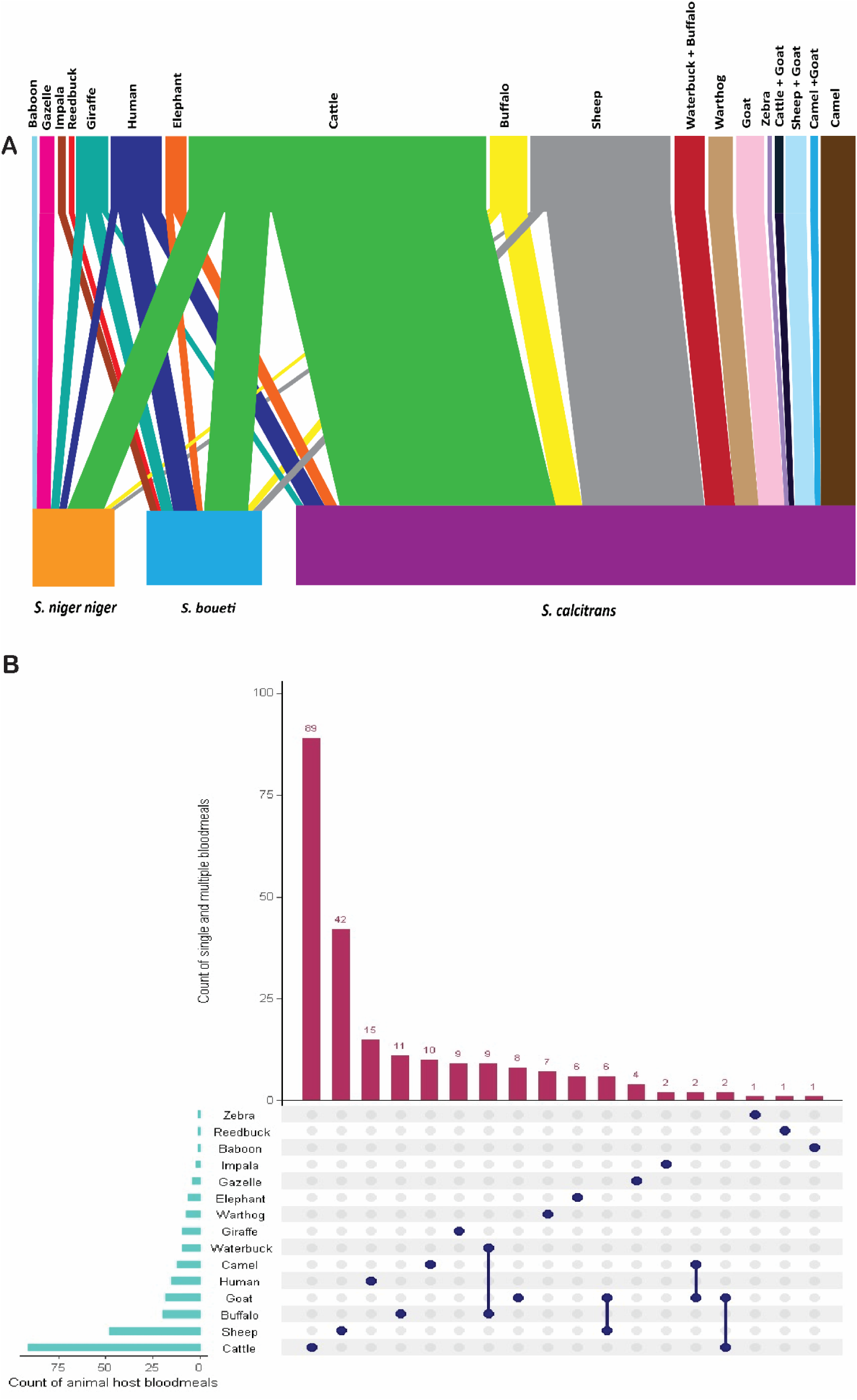
Identification of vertebrate hosts from bloodmeal analysis of *Stomoxys* spp. (A) A *bipartite* network graph showing feeding interactions between hosts and blood-fed *Stomoxys* spp. The top bar indicates hosts while the bottom bar indicates the *Stomoxys* spp. while the lines illustrate the interaction. The size of a bar reflects the number of blood-fed *Stomoxys* (if it is a bottom bar) or the number of mammalian hosts that were fed on the vector (if it is a top bar). The thickness of a line corresponds to the number of blood-fed hosts detected in the various *Stomoxys* spp. (B) An *Upset* plot showing the total number of hosts fed per species and also multiple host feeding.

### 3.3 *Stomoxys* flies and domestic animals harbor various hemopathogens

Another data required to elucidate the role of *Stomoxys* for various pathogen transmission dynamics besides blood meal source is to study the pathogens network between *Stomoxys* and some of the most preferred host animals they feed on. Various pathogens were detected both in the blood of livestock which were also common in the *Stomoxys* flies. *Anaplasma* sp., *Theileria* sp., and *Trypanosoma* sp. were shared across all analyzed domestic animal hosts and *Stomoxys* flies. *Ehrlichia* sp. and *Rickettsiae* sp. were detected in both camels and cattle. *Coxiella burnetti* was only detected in camels. In camels (n= 452), *Anaplasma* sp., was the most prevalent pathogen affecting 64.7% of the camels. *Trypanosoma* sp. was detected in 12.3% and *Ehrlichia* sp. in 12.2% of camels sampled. *Coxiella burnetti* was found in 6% of the camels, while *Rickettsiae* sp. was the least detected in 4% of the camels. We did not detect any *Theileria/Babesia* spp. in camels. Additionally, in cattle out of n=124, we found a high prevalence of *Theileria*/*Babesia* sp. with 56.6% and *Anaplasma* sp. in 54.1%. *Trypanosoma* sp. was detected in 10% of cattle, *Rickettsiae* sp. in 5.7% of the cattle and *Ehrlichia* sp. was the least prevalent, found in only 1.6% of cattle. Among all hosts, sheep had the least pathogen diversity with only *Theileria/Babesia* sp. being detected in 4% of the sheep. A total of 3,451 *Stomoxys* were screened for pathogen diversity across the study counties. Among these, *Anaplasma* sp., was the most frequently detected with 49.1%, 19.1% having *Theileria/Babesia* sp. and 9.1% having *Trypanosoma* sp. For comparison, *Glossina pallidipes* (n=1000) co-inhabit with *Stomoxys* had pathogen prevalence of *Trypanosoma* sp., *Anaplasma* sp., and *Theileria/Babesia* sp., which were detected in 7.5%, 4%, and 11% of the flies, respectively.

### 3.4 *Stomoxys* flies are competent mechanical vectors of *T. evansi* and *T. vivax*

*Stomoxys* feeds about 9.98 ± (5.5) mL blood when fully engorged and needs on average 4.6 ± (2) minutes to fully engorged, the number in parenthesis is the standard deviation of the mean (n=10 flies). *T. evansi* survived in various tissues of *Stomoxys* after immediate disruption of feeding. About 30% of *Stomoxys* fed on infected mice showed parasites in the proboscis if feeding was interrupted within one minute. However, more than 80% of the flies fed on infected mice had parasites in their crops and gut when feeding was interrupted within one minute. *T. evansi* survived up to 5 hours in the gut of *Stomoxys* which shows a possibility of delayed transmission of trypanosomiasis. In the first three hours, the trypanosomes were very active swimmers, and gradually became inactive after 4 hours and were all dead 6 hours post-feeding by flies. We demonstrate that *Stomoxys* flies transmit *T. evansi* through *in vivo* experiments using laboratory mice, with 8.3% (2/ 24) mice with patent parasitemia detected by microscopy by day 7 after infection assays. The wild *T. evansi* strain showed moderate virulence as the mice maintained a peak of parasitemia (1 × 10^8^ trypanosomes/ml blood) for several days and died between the 10^th^ and 14^th^ days, respectively with mild clinical symptoms. Concerning *T. vivax,* we found longer survival times in *Stomoxys* guts as compared to *T. evansi,* as we found live *T. vivax* at 16 hours, as opposed to 6 hours for *T. evansi* (Fig. 6). Furthermore, we found a higher transmission success rate as compared to *T. evansi*, as we showed 30% (3/10) transmission was successful. The incubation period varied from 6 days to 11 to 34 days in the three *T. vivax*-infected mice. The *T. vivax* IL 2136 strain exhibited a moderate level of virulence as the infected mice also sustained a high parasitemia of 1 × 10^8^ trypanosomes/ml blood for several days and died on the 6^th^, 8^th^, and 12^th^ days. For comparison, we did the same mechanical infection experiment with *G.pallidipes* with 5 trails and all transmitted *T.evansi* to five mice, demonstrating variation between *Stomoxys spp*. and *G.pallidipes*.

### 3.5 Natural pathogen transmission assays through feeding bites on experimental mice by field-collected *Stomoxys* spp

We finally asked if field-collected *Stomoxys* flies are capable of transmitting pathogens they harbored by allowing field-trapped *Stomoxys* flies to feed on healthy mice. We demonstrated wild caught *Stomoxys* flies are capable of delayed transmission of various pathogens they harbored in *in vivo* experiments*. Stomoxys* flies transmitted *Theileria mutans* (GenBank Accession Number, PP918990) into healthy mice after delayed feeding in the field (Fig. 5B) Furthermore, all mice showed *Anaplasma spp*. infection microscopically only. These pathogens had low virulence as the mice showed no clinical symptoms and no mortality of the mice was recorded for > 120 days.

**Figure 5:**
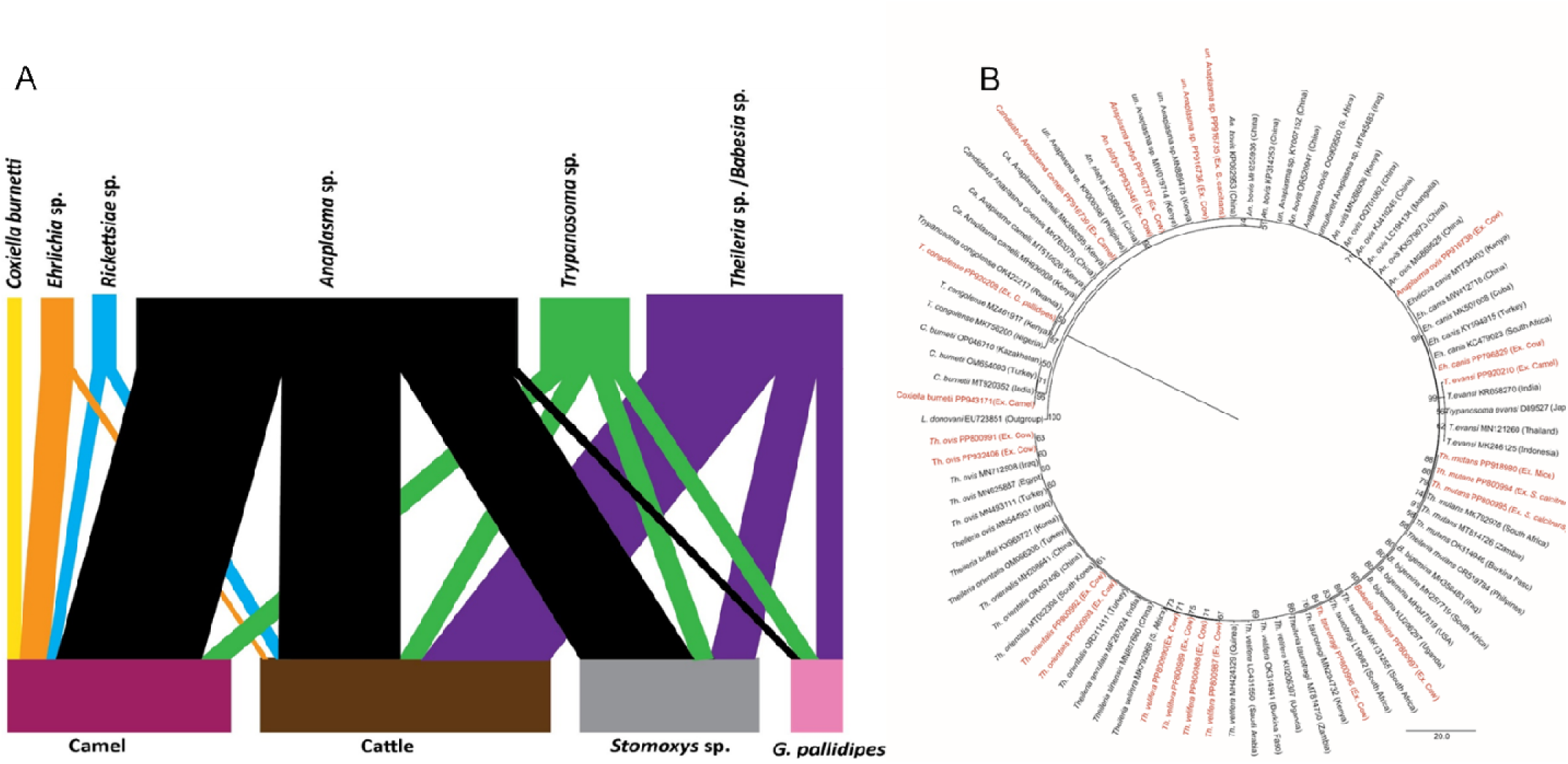
Pathogen diversity in host animals and vectors from various sites through molecular screening and neighbor-joining tree showing pathogens. (A) A bipartite network graph showing pathogen interactions between hosts (camel, cattle, and sheep) and vectors *Stomoxys* sp. and *Glossina* sp. The top bar indicates pathogens while the bottom bar indicates the hosts and vectors while the lines illustrate the interaction. The thickness of a line corresponds to the number of pathogens detected in either the hosts or vectors. (B) Neighbor-joining tree showing pathogens from host animals and vectors.

**Figure 6:**
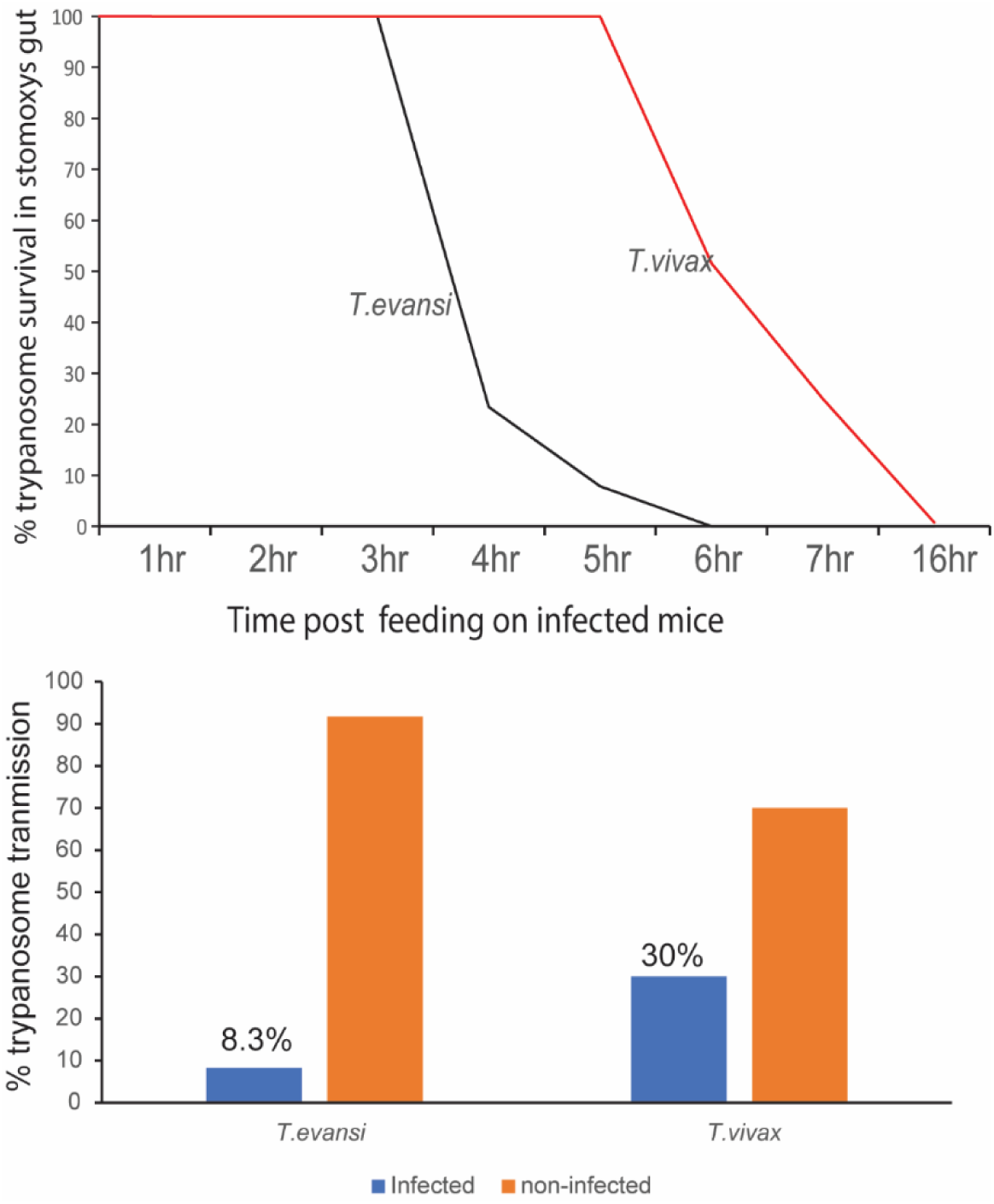
Vector competence of *Stomoxys* spp. to transmit trypanosomes. (A). Graph showing the survival of *T. evansi* and *T. vivax* in *Stomoxys* gut (B). The success of infection.

## 4. Discussion

In this study we aim to understand *Stomoxys*-host-pathogens network interaction to get insight about the role of *Stomoxys* flies in disease transmission dynamics, and how transmission networks of pathogens-vectors-host are functioning. Out of 18 species of *Stomoxys* that are found globally 14 of them are found in Africa [1]. We found year-round wide distribution of six species of *Stomoxys* including *Stomoxys calcitrans, S. sitiens, S. niger niger, S. niger bilineatus*, *S. boueti* and *S. taeniatus* that varies in their abundance and diversity in nine regions including in three tsetse infested ecologies. With our wider geographic coverage, we reported only six species of *Stomoxys* as compared to Mihok et al., 1996 [33] who did trapping from Nairobi National Park and reported ten species of *Stomoxys*. *Stomoxys* species complexity varies between ecologies, national reserve got more species of *Stomoxys* as compared to zero grazing ecologies. For example, species diversity was notably high in Kwale County, most likely due to the availability of numerous breeding sites generated by the forested terrain [34]. Kiambu County, where zero grazing is implemented, had a high population density of mainly three species, which is due to the availability of readily available breeding substrate [35]. Isiolo (n = 2,514, 22.20%), Kajiado (n= 1,373, 12.12%), and, Homabay (n = 545, 4.81%) counties exhibited a considerably high population of *Stomoxys*, which could be attributed to the habitat, which has a semi-arid climate that encourages the growth of the flies (Mavoungou et al., 2017). Marsabit (n = 74, 0.65%) and Samburu (n = 26, 0.23%) counties had the least abundance, which could be attributed to the environment, which is a hot and arid climate that is not friendly to *Stomoxys* because high temperatures have been reported to cause a drop in the fly population due to reduced survival of larvae and pupae [36].

To get insight of the role of *Stomoxys* in disease transmission dynamics we need to understand the natural feeding habits of *Stomoxys* flies from various ecologies. We showed *Stomoxys* flies feed on a wide range of wild and domestic animals, including humans which is comparable to tsetse flies [37], [38] and which also corresponds to prior research findings [2], [3], [39]. From our study *Stomoxys* need an average of 4 minutes to complete feeding, this may induce host defense and interrupted feeding, which will result in multiple hosts feeding and pathogen transmission [40]. This disruption of *Stomoxys*-feeding before bloodmeal completion enables the vectors to switch to new hosts to continue feeding, which serves as the basis of mechanical transmission of pathogens [5]. In general, the relatively wide variety of feeding suggests that *Stomoxys* may take a more opportunistic approach to host selection, potentially responding to the availability of susceptible hosts in its environment [39]. We found 225 blood-fed *Stomoxys* out of 3451 showing that most *Stomoxys* flies caught using traps are often seeking hosts for a blood meal, which is also true for other hematophagous insects [41]. Moreover, blood digestion starts more rapidly in *Stomoxys* as compared to other hematophagous flies [42]. Thus, the low rate of blood meal identifications could be explained by the degradation of host DNA during digestion in the fly midgut or furthermore, *Stomoxys* takes too little blood quantity in the midgut, even in strained animals *Stomoxys* takes only 10 mg in 4 minutes. Nevertheless, the diversity of hosts we successfully identified includes diverse wild, domestic animals and humans (*Homo sapiens*). The diversity of blood meals can be due to the flies high mobility, their opportunistic feeding behavior, and their frequent feeding habit. Furthermore, trap position and ecologies may influence the range of host species *Stomoxys* may feed on. For instance, Mavoungou et al., 2008 demonstrated that *Stomoxys* flies sampled in canopies mainly feed on arboreal species [43]. We can also notice the absence of small mammals (e.g., rodents) within the diversity of host vertebrates we identified. This may be explained by the trophic preferences of *Stomoxys* flies, the same as tsetse for large vertebrates [37], [44], [45].

Such a diverse feeding host will expose *Stomoxys* to diverse pathogens as the host varies in their pathogen reservoir capacity [46], which is shown in our pathogen network result. The epidemiology of African trypanosomiasis includes the biting rate of vectors on infected hosts and the probability of vectors feeding on different hosts as key parameters for understanding the transmission of these infections. Molecular pathogen screening led to the identification of various pathogens that showed epidemiological overlap and interactions between hosts and vectors in the study area. Concerning the pathogens network, we detected high infection rates of *Anaplasma* spp. both in selected domestic animals and *Stomoxys*. The detection of the pathogen from the biting flies confirms the possibility of these flies acquiring and maintaining these pathogens. Ticks including *Rhipicephalus decolaratus, R. microplus*, *Hyalomma marginatum rufipes, R. evertsi*, and *R. simus* are among the *Anaplasma* biological vectors [47]. However, Scoles et al. (2005) found that stable flies can transmit *A. marginale* [48] whereby they demonstrated that the Florida strain of *A. marginale* which cannot be transmitted by ticks, was more effectively maintained in stable fly mouth parts compared to the tick-transmittable St. Maries strain [5]. Similarly, Bargul et al. (2021) demonstrated *A. camelii* transmission by *Hippobosca camelina*, but the same pathogens were not detected in ticks collected from *A. camelii-*infected camel [49]. According to a report conducted by Oliveira et al., 2011, seroprevalence and the presence of tabanids and stable flies are associated with bovine exposure to *A. marginale*, which is widespread in Costa Rican dairy herds [50]. Another pathogen found with high prevalence both in the host and *Stomoxys* was *Theileria* spp. and our *in vivo* experiment demonstrated the successful transmission of *Theileria mutans* by *Stomoxys* flies. Similarly, *Theileria* DNA was detected in stable flies, in the case of *T. orientalis* at least for two hours after blood-feeding in a study done by [51]. Interestingly, we did not detect anyof *Ehrlichia* spp. and *Rickettsia* spp. in *Stomoxys* or *Glossina* but the pathogens were present in both hosts, demonstrating the poor vector competence of *Stomoxys* for these particular pathogens.

In our *in vivo* experimental studies, we discovered that *T. evansi* could actively persist in several tissues of *Stomoxys* flies. *Stomoxys* flies displayed motile *T. evansi* in the proboscis after immediate feeding disruption, which could be observed for up to 5 minutes. These findings imply that the mouthparts of *Stomoxys* species do not promote trypanosome survival for long [12]. This may be due to the direct transit of blood to the midgut during eating, which leaves very little blood in the proboscis [52]. Our findings are consistent with those of Sumba et al., 1998, who confirmed that motile and presumably viable trypanosomes remained in or on the proboscis for around 5-7 minutes after feeding was terminated [12]. While in the midgut, we established *T. evansi* could survive for up to 5 hours in the gut of *Stomoxys*. This survival capability allows *T. evansi* to allow for a second possible mechanism of transmission, namely regurgitation [13]. Reports in other literature summarized different survival times for various trypanosome species in different biting flies. For instance, Sumba et al. 1998 found that *T. congolense* could live up to 3 and half hours and *T. evansi* up to 8 hours in the guts of *S. niger* and *S. taeniatus*. Additionally, Getahun et al. 2022 found that *T. congolense* could live for 3 hours and trypanozoons for 5 hours in the midgut of *S. calcitrans* [3], [12], [13]. Additionally, *Stomoxys* flies are efficient mechanical vectors of *T.vivax.* We showed that *T.vivax* survived in the *Stomoxys* gut for a longer period as compared to *T.evansi* for unknown reasons. The experimental assay showed that *in vivo* transmission of *T. evansi* and *T.vivax* by *Stomoxys* flies was successful with variable success rates. Our findings align with that of Mihok et al., 1995 where it was established *S. calcitrans* transmits multiple trypanosome species with various transmission rates. A contrary finding by [53] using relevant host cattle-*T. vivax*-and *S. calcitrans* interaction reported that *S. calcitrans* could not transmit *T. vivax* to cattle this could be due to the transmission experiment design. The authors released the flies into a pen with both healthy and infected animals, flies may be more attracted to an infected host than a healthy one [54] in our protocol establishment when both the infected mice and healthy mice were kept together we found no transmission, despite 20 trials. Furthermore, the interrupting feeding was done after 1.5 minutes, in our protocol establishment experiment when flies were allowed to feed for more than 1 minute, and transferred to a new host, they lost motivation to feed immediately and even those that fed later did not transmit, it seems mechanical transmission of trypanosomes is time sensitive. Furthermore, mechanical transmission is parasitemia dependent [30] we found the best parasitemia range for mechanical transmission was 1 × 10^8^ trypanosomes/mL blood required The other factor could be *Stomoxys* species may vary in their vector competence, in our experiment we kept the *Stomoxys* species complex intact mainly composed of three species as a matrix.

## 5. Conclusion

The wider geographic distribution, fast reproduction, species diversity, year all presence, diverse feeding habits, the plethora of pathogens harbored, and their successful vectorial capacity of transmitting *T. evansi, T. vivax, Anaplasma* spp., and *Theileria mutans* as shown by our *in vivo* experiments demonstrate *Stomoxys* flies are significant but overlooked vectors of various pathogens of livestock. *Stomoxys* flies may play a significant role in the spread and maintenance of *T. evansi* and *T. vivax* in the wide geographic regions of the world. In the future, it is important to do vector competence experiments using a specific *Stomoxys* spp. with relevant host animals -pathogens interaction. In our experiment we kept the natural species complex, composed of mainly *S. calcitrans S. niger niger,* and *S. boueti* matrix intact, which means we did not try to separate them by species, in the future it is important to do individual species vector competence.

## Data availability

All relevant data are in the manuscript and supplementary data. All sequences have been deposited in the NCBI database

## Author’s contributions

J.W.M; Conceptualization, Data curation, Formal analysis, Investigation, Methodology, Visualization, Writing – original draft, Writing – review & editing. J.L.B; Conceptualization, supervision, Writing-review & editing. J.M.O.M; Data curation, Methodology, Formal analysis, Writing-review & editing. E.M.N; Data curation, Methodology, Formal analysis, Writing-review & editing. S.K.T; Data curation, Methodology, Formal analysis, Writing-review & editing. D.K.M; Conceptualization, Funding acquisition, Resources, Supervision, and, Writing-review & editing. M.N.G; Conceptualization, Funding acquisition, Resources, Supervision, Writing – original draft, Writing-review & editing, and Investigation. All authors read and commented on the content.

## Acknowledgments

We would like to thank Dr. Geoffrey Gimonneau and Dr. Marc Desquesnes for useful discussion about infection experiment protocol development. We acknowledge Dr. Steve Mihok for the useful discussion and his support in Stomoxys identification. James Kabii for his technical support; John Ngiela, Victor Omondi, and Peter Ahuya helped. We are grateful to Shadrack Kibet for designing the map of sampling sites. Joseck Esikuri for supplying mice for experimental pathogen transmission assays. Caroline Muya helped in handling the administrative aspects relating to this study.

